# HPV16 E2 protein possesses intrinsic helicase activity and sterically hinders E1 function through direct interaction

**DOI:** 10.1101/2025.09.15.676238

**Authors:** Ping Xu, Shuning Cai, Lihong Zhang, Yueqi Wu, Ke Xu, Yigang Tong, Shan Xu

**Affiliations:** BAICSM, State Key Laboratory of Green Biomanufacturing, College of Life Science and Technology, Beijing University of Chemical Technology, Beijing 100029, China; Department of Pharmacy, The Second Qilu Hospital of Shandong University, Jinan 250033, China

**Keywords:** HPV16, E2, E1, helicase, ATP

## Abstract

HPV16 E2 protein is a key regulatory protein essential for viral replication, yet no enzymatic activity had been attributed to it until now. In this study, we report for the first time that E2 possesses intrinsic ATP-dependent DNA unwinding activity. Mutational analysis identified residues K299, Y303, and K306 as critical for this helicase function. We further demonstrate that podophyllotoxin directly binds to E2 and inhibits its unwinding activity with an IC_50_ of 0.1074 µM, mediated primarily by residues Q320 and H342. Comparative analysis revealed that the ATPase and helicase activities of E2 are considerably weaker than those of E1. Notably, we discovered that E2 potently inhibits the helicase activity of E1. This suppression is facilitated by the N-terminal domain of E2 (amino acids 1–245) through direct interaction with E1, with residue E39 playing a critical role. Our findings not only unveil a previously unrecognized enzymatic function of E2 but also suggest its role as a potential antiviral target. Moreover, the observed inhibitory effect of E2 on E1 highlights a novel regulatory mechanism for HPV DNA replication.

**Synopsis:** 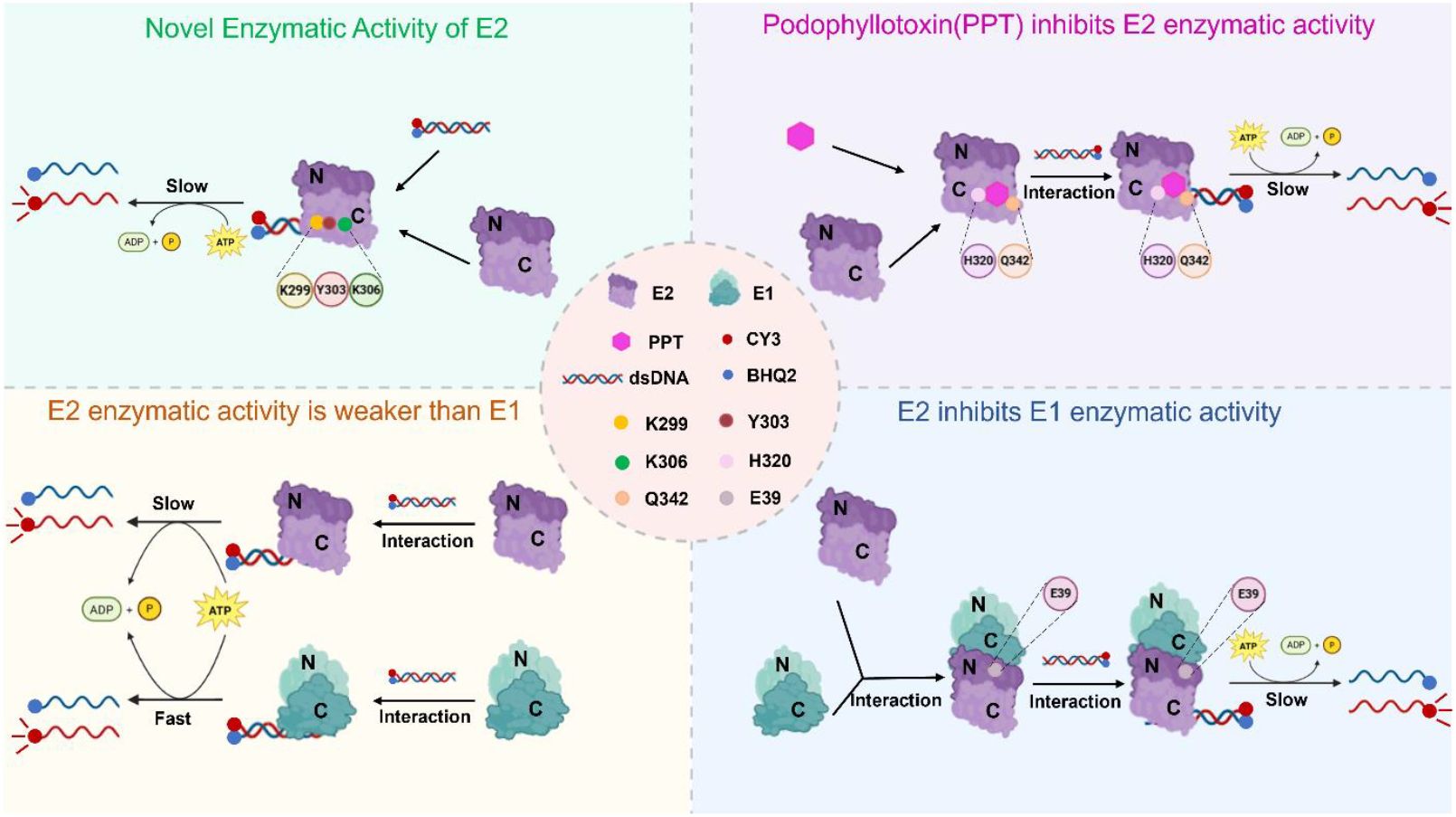

E2 protein has traditionally been recognized primarily for its DNA-binding and transcriptional regulatory functions. This study provides the first evidence that E2 protein possesses intrinsic enzymatic activity, identifies a small-molecule inhibitor targeting this activity, and reveals a novel mechanism of E1-E2 interaction.

- HPV16 E2 Protein Exhibits ATPase and Helicase Activities
- Identification of Key Amino Acid Residues for HPV16 E2 Protein Enzymatic Activity
- PPT Effectively Inhibits E2 Protein Helicase Activity *In Vitro*
- E2 Protein Exhibits Weaker Enzymatic Activity Than E1 and Inhibits E1 Helicase Activity
- E2 Inhibits E1 Helicase Activity Through Protein-Protein Interaction

## Introduction

Human papillomavirus (HPV) is one of the most common sexually transmitted infections worldwide. Recent epidemiological data from the World Health Organization (WHO) indicate that approximately 11–15% of reproductive-aged women (15–49 years) harbor genital HPV infections, of which 4–7% are attributed to high-risk genotypes, constituting a substantial public health burden (Bruni et al,2023). HPV-associated malignancies account for an estimated 342,000 annual deaths globally, with cervical cancer representing 88% of this burden, disproportionately affecting low- and middle-income countries (Deng et al,2004;Sung et al,2021; Bray et al,2024; Brisson et al, 2020; Drolet et al, 2021).This data highlights the significant impact of HPV infection on health equity, particularly in regions with limited healthcare resources. HPV is a non-enveloped double-stranded DNA virus that specifically infects cutaneous and mucosal epithelial tissues (McBride,2022; Arbyn et al, 2020; Pešut et al,2021; Doorbar et al,2015; Doorbar et al,2012). More than 130 genotypes have been characterized, each exhibiting distinct pathological manifestations ranging from benign lesions (e.g., common warts, condylomata acuminata) to malignant transformations (Serrano et al,2012; D’Souza et al,2023; Haley et al,2019; Muñoz et al,2003; Al-Awadhi et al,2019). High-risk genotypes, notably HPV16 and HPV18, are established etiological agents of anogenital and oropharyngeal carcinomas (Koster et al,2022; Arroyo et al,2024; Whop et al,2019; Billingsley et al,2022).

HPV 16, a predominant high-risk variant, is intimately associated with oncogenesis, particularly cervical carcinogenesis (Schiffman et al,2007; Meng et al,2024; Nelson et al,2023). The viral genome comprises an approximately 8-kb circular double-stranded DNA encoding eight canonical open reading frames (E1, E2, E4, E5, E6, E7, L1, L2). These genes are functionally partitioned into early regulatory proteins (E1–E7), governing viral replication, transcription, and cellular transformation, and late structural proteins (L1, L2), constituting the viral capsid. The early proteins E1 and E2 orchestrate pivotal stages of the viral life cycle, including initiation of DNA replication, transcriptional regulation, and virion morphogenesis (Hebner et al,2006). E1 functions as a multifunctional helicase that recognizes the viral origin of replication (ori) and recruits host replication factors (Sakakibara et al,2011; Nakahara et al,2015; Schuck et al,2011). Conventional models posit E2 as a sequence-specific DNA-binding protein modulating transcriptional activation and suppression, while also stabilizing E1 and facilitating its nuclear localization (Arbyn et al, 2020; Hou et al,2002; McBride,2013). Furthermore, E2 augments E1-mediated replication through direct protein–protein interactions (Yilmaz et al,2023; Sedman et al,1998; Longworth et al,2004; Chen et al,2002). Nevertheless, the autonomous enzymatic potential of E2 remains unexplored, and the precise stoichiometry and dynamics of E1–E2 complexes warrant further elucidation.

In this study, we developed a real-time duplex DNA unwinding assay leveraging Fluorescence Resonance Energy Transfer (FRET) to quantitatively characterize the helicase activity of HPV16 E1 and E2. We demonstrate that HPV16 E2 exhibits intrinsic ATP-dependent DNA unwinding activity, a function previously unassociated with this viral factor. The key residues at the ATP-binding site (Y303) and DNA-binding site (K299, Y303, and K306) of E2 are crucial for its ATPase and helicase activities. Mutations at these sites significantly impair both the ATPase and helicase activities of the E2 protein. Moreover, we identified podophyllotoxin (PPT) as a potent inhibitor of E2 helicase activity, with half-maximal inhibitory concentration (IC_50_) values in the nanomolar range. Crucially, E2 markedly attenuates E1-mediated unwinding kinetics in an interaction-dependent manner. Disruption of E2–E1 binding—achieved through interface mutations—abolished this trans-regulatory effect. Our work provides the first evidence of catalytic functionality in HPV16 E2, unveils a novel regulatory mechanism governing E1–E2 cooperativity, and identifies a pharmacologically actionable site for antiviral development. These insights advance fundamental understanding of HPV replication mechanics and offer new avenues for therapeutic intervention targeting viral helicase complexes.

## Results

### HPV16 E2 Protein Exhibits ATPase and Helicase Activities

The HPV E2 protein has long been considered an auxiliary protein, and its intrinsic enzymatic activities remain to be fully characterized. To investigate the ATPase and helicase activities of HPV16 E2 protein, the full-length E2 protein (E2, residues 1–365) was expressed and purified from *Escherichia coli*. Coomassie blue staining and Western blot analysis confirmed that the molecular weights of these proteins were correct and consistent with theoretical calculations (Figures 1A and 1B). Helicases typically unwind nucleic acids using energy derived from ATP hydrolysis. We measured the ATPase activity of the E2 protein using a malachite green-based colorimetric assay, which detects free inorganic phosphate released during ATP hydrolysis. The data were fitted according to the Michaelis-Menten equation and presented as a double-reciprocal plot (Figure 1C). The results demonstrated that the purified full-length HPV16 E2 protein possesses ATP hydrolysis activity (Vmax=0.52 μmol/l·min,Km=0.11 mmol/l). Next, the degree of DNA strand separation by HPV16 E2 protein was monitored in real time by FRET technology (Figure 1D). Complementary nucleic acid strands were labeled with the fluorophore Cy3 or the quencher BHQ2. When the complementary strands annealed into a duplex, the fluorescence emitted by Cy3 was largely absorbed by BHQ2, resulting in a low Cy3 fluorescence signal. As the helicase unwound the duplex, the quenching effect of BHQ2 on Cy3 decreased or disappeared, leading to a corresponding increase in Cy3 fluorescence. As shown in Figure 1D, HPV16 E2 protein exhibits ATP hydrolysis-dependent DNA unwinding activity.

**Figure 1.**
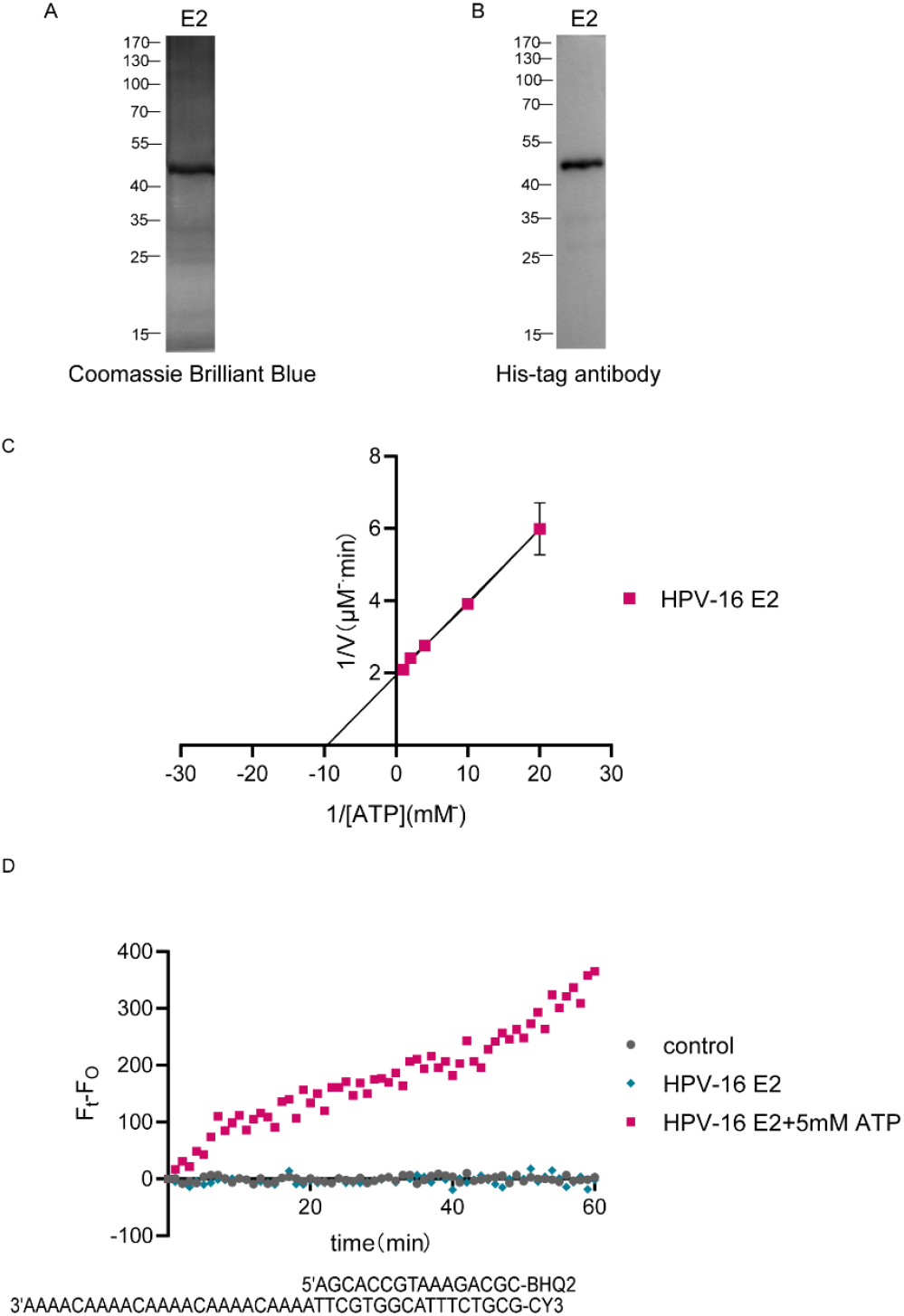
Purification and enzymatic activity assays of HPV16 E2 protein. **(A)** SDS-PAGE analysis of purified E2 protein stained with Coomassie blue. **(B)** Western blot analysis of purified E2 protein using an anti-His antibody. **(C)** ATPase activity of purified E2 protein. ATP hydrolysis assays were performed with 2.5 μM protein in the presence of the indicated ATP concentrations at 37°C for 60 min. Data were fitted to the Michaelis-Menten equation and presented as a double-reciprocal plot. Error bars represent the standard error of replicate measurements. **(D)** Real-time unwinding kinetics of 0.4 μM E2 protein on a fluorescently labeled 16-bp dsDNA. Buffer without protein was used as a control. Helicase activity of E2 with or without ATP were detected. Fluorescence signals were measured every 1 min. The unwinding extent was defined as F_t_−F_0_, where F_t_ is the fluorescence of the sample at a given time and F_0_ is the initial fluorescence of the sample. The sequence information of the dsDNA is shown at the bottom of the corresponding assay plot.

### Identification of Key Amino Acid Residues for HPV16 E2 Protein Enzymatic Activity

Molecular docking was performed between the E2 protein and ATP, followed by PyMOL visualization to identify interacting amino acid residues. Among these, residue Y303 of the E2 protein formed a hydrogen bond with ATP (Figure 2A). This site was subsequently mutated, and Western blot analysis confirmed that the mutant protein exhibited the correct molecular weight, consistent with theoretical calculations (Figure 2B). ATP hydrolysis assays revealed that the Y303N mutation significantly impaired the ATPase activity of E2 (Figure 2C), indicating that Y303 is a critical residue for E2’s ATPase function. The E2 protein typically functions as a dimer when binding to DNA. Molecular docking of the dimeric E2 protein with DNA, followed by PyMOL visualization, identified additional interacting residues—K299, Y303, and K306—which formed hydrogen bonds with DNA (Figure 2D). These sites were mutated, and Western blot analysis confirmed that the resulting mutant proteins had the expected molecular weights (Figure 2E). Among different HPV genotypes, the catalytic residues K299 and Y303 are highly conserved, whereas K306 is not (Supplementary Figure S2). When the K299R, Y303N, and K306A mutants were subjected to dsDNA unwinding assays, they exhibited significantly reduced helicase activity and slower unwinding rates (Figure 2F). We also measured the ATPase activity of the K299R and K306A mutants and found that the mutations at these sites did not affect their ability to hydrolyze ATP (Supplementary Figure S6). Additionally, we tested the N296A mutant in the dsDNA unwinding assay and observed similarly diminished helicase activity and slower unwinding kinetics, and this residue is highly conserved across different HPV genotypes (Supplementary Figures S2 and S4). We speculate that this residue, located near the helicase active region, may indirectly affect enzymatic function. These results demonstrate that these four amino acids are critical for the helicase activity of the E2 protein and further confirm that the enzymatic domain of E2 resides in its C-terminal region (residues 245–365).

**Figure 2.**
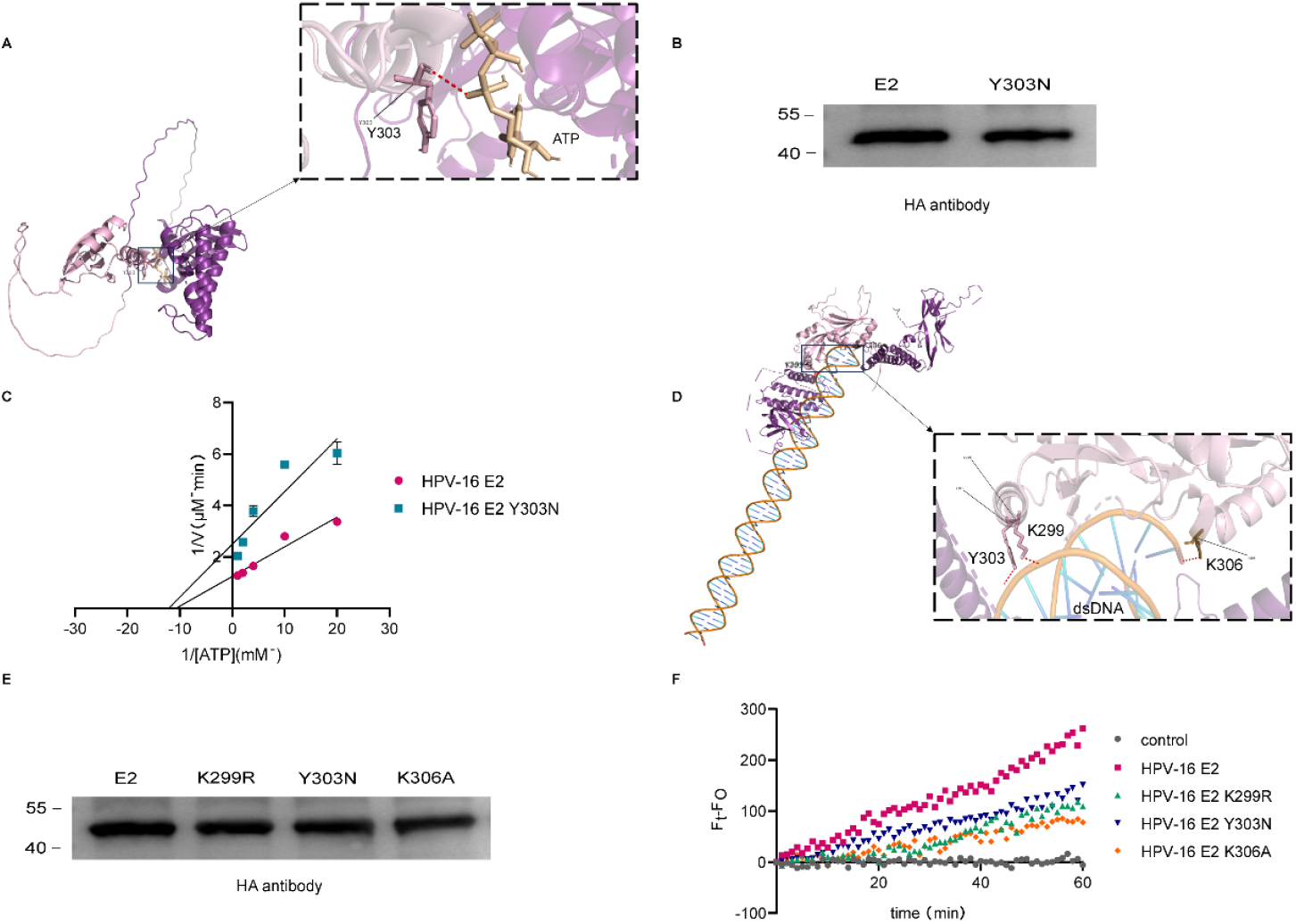
Identification of key amino acid residues for HPV16 E2 protein enzymatic activity. **(A)** Molecular docking of E2 protein with ATP using AutoDock, showing hydrogen bond formation between Y303 and ATP. ATP is shown in dark yellow, Y303 in pink, and hydrogen bonds in red. **(B)** Western blot analysis of purified wild-type E2 and mutant Y303N proteins using an anti-HA antibody. **(C)** Comparison of ATPase activity between 2.5 μM wild-type E2 and mutant E2 Y303N. **(D)** HDOCK-based docking of dimeric E2 protein with DNA, revealing hydrogen bonds between K299, Y303, K306, and DNA. K299 and Y303 are shown in pink, while K306 from the other monomer is shown in dark brown. DNA is represented in dark yellow, and hydrogen bonds are indicated in red. **(E)** Western blot analysis of purified E2 protein and mutants K299R, Y303N, and K306A using an anti-HA antibody. **(F)** Real-time unwinding kinetics of 0.4 μM wild-type E2 and mutants K299R, Y303N, and K306A on fluorescently labeled 16-bp dsDNA. Buffer without protein served as a control. Fluorescence signals were measured every 1 minute. Unwinding extent was defined as F_t_−F_0_, where F_t_ is the sample fluorescence at a given time and F_0_ is the initial fluorescence.

### Podophyllotoxin (PPT) Effectively Inhibits E2 Protein Helicase Activity *In Vitro*

Podophyllotoxin (PPT) is a natural product currently used to treat external warts caused by HPV-6 and HPV-11(Kollipara et al,2015; Komericki et al, 2011; Gilson et al,2020; Murray et al,2018; Nicolaidou et al,2021). However, its mechanism of action remains unclear. Molecular docking between the E2 protein and podophyllotoxin suggested their potential interaction. Residue H320 and Q342 of the E2 protein formed a hydrogen bond with PPT (Figure 3A). Bio-Layer Interferometry (BLI) was used to further validate this interaction, confirming that they bind to each other with an affinity (KD) of 6.234 × 10^−4^ M (Figure 3B). To investigate whether PPT inhibits the helicase activity of HPV16 E2 in vitro, a FRET-based assay was employed (Figure 3C). The inhibitory effect of PPT on E2 helicase activity was characterized by measuring the extent of dsDNA unwinding by E2 at increasing concentrations of PPT. The half-maximal inhibitory concentration (IC_50_) was determined to be 0.1074 µM (Figure 3D), indicating that PPT effectively inhibits the helicase activity of E2 *in vitro*. The conserved residues H320 and Q342 were mutated to alanine (Supplementary Figure S2),and the resulting mutant proteins were verified by Western blotting, which confirmed that their molecular weights matched the theoretical values (Figure 3E). When these two residues were mutated, the inhibitory effect of PPT on helicase activity was abolished (Figure 3F), suggesting that H320 and Q342 are critical for the interaction between E2 and PPT. In conclusion, PPT inhibits the enzymatic activity of E2 by directly binding to the protein, establishing it as a small-molecule inhibitor of the E2 helicase.

**Figure 3:**
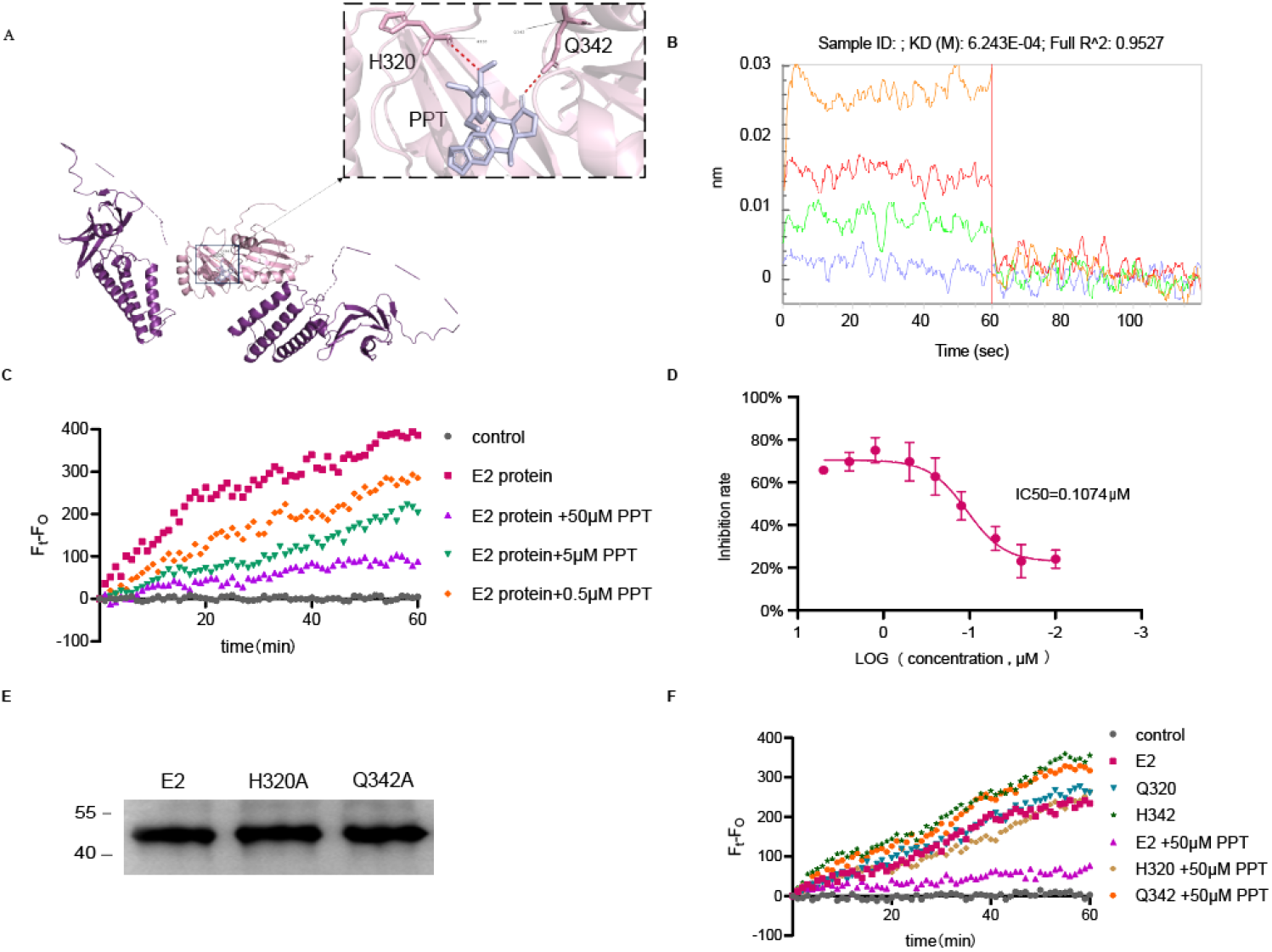
Podophyllotoxin (PPT) effectively inhibits E2 protein helicase activity *in vitro*. **(A)** Molecular docking of E2 protein with PPT using AutoDock, showing hydrogen bond formation between H320、Q342 and PPT. PPT is shown in light purple, H320、Q342 in pink, and hydrogen bonds in red. **(B)** Bio-Layer Interferometry (BLI) analysis of the interaction between E2 protein and PPT. The curves from bottom to top represent working concentrations of PPT at 50, 100, 200, and 400 μM, respectively. Data were processed using double-referenced subtraction (sample and sensor reference) based on a 1:1 binding model. Kinetic parameters (kon and koff) and the affinity constant (KD) were obtained using Octet BLI Analysis 12.2 software. **(C)** Real-time FRET-based detection of the inhibitory effect of PPT (0.5 μM, 5 μM, 50 μM) on the unwinding activity of 0.4 μM E2 protein. **(D)** Determination of the IC50 value for PPT. Using the FRET assay, the reaction mixture was added to a white 96-well plate and covered with foil. After 1 hour of incubation, fluorescence was measured using a microplate reader with excitation at 550 nm, emission at 620 nm, and a cutoff at 610 nm. The assay was performed with 0.4 μM E2 protein and PPT concentrations ranging from 0.1 mM to 0.00025 mM (0.1, 0.05, 0.025, 0.01, 0.005, 0.0025, 0.001, 0.0005, and 0.00025 mM). The experiment was repeated three times, and the IC50 was calculated using GraphPad Prism 8 software. The graph displays the mean ± standard deviation of three independent experiments. **(E)** Western blot analysis of purified wild-type E2 and mutant H320A, Q342A proteins using an anti-HA antibody. **(F)** Helicase assays measuring dsDNA unwinding by wild-type E2 or mutants H320A and Q342A in the presence or absence of PPT. Reactions contained 0.4 μM wild-type or mutant E2 proteins and 50 μM PPT, with 16-bp dsDNA as substrate.

### E2 Protein Exhibits Weaker Enzymatic Activity Than E1 and Inhibits E1 Helicase Activity

Purified HPV16 E1 protein was confirmed by Coomassie blue staining and Western blot analysis to have the correct molecular weight, consistent with theoretical calculations (Figure 4A). As previously established, HPV16 E1 protein is an ATP-dependent helicase (Baedyananda et al,2022; Orav et al,2019; Bandyopadhyay et al,2008; Hughes et al,1993; Rana et al,2025; Deng et al, 2004). We compared the ATPase activities of E1 and E2 proteins using a malachite green-based colorimetric assay that detects free inorganic phosphate released during ATP hydrolysis. Data were fitted to the Michaelis-Menten equation and presented as double-reciprocal plots (Figure 4B), revealing significantly weaker ATPase activity in E2 compared to E1 (E2: Vmax=0.34 μmol/L·min, Km=0.06 mmol/L; E1: Vmax=0.84 μmol/L·min, Km=0.14 mmol/L).FRET-based assays comparing helicase activities showed that E2 unwound dsDNA at a markedly slower rate than E1, confirming its weaker helicase activity (Figure 4C). Since E2 typically functions as an auxiliary protein cooperating with E1 in viral DNA replication regulation, we investigated whether E2 affects E1 helicase activity. Surprisingly, E2 was found to inhibit E1 helicase activity (Figure 4D). However, malachite green assays demonstrated that E2 does not affect helicase activity of E1 (Figure 4E). We hypothesize that the N-terminal domain of the E2 protein (residues 1–245) binds to the C-terminal region of the E1 helicase (residues 143 – 645), thereby obstructing its ability to bind the dsDNA substrate. Consequently, the E2 protein assumes the helicase function, binding to dsDNA and unwinding the double strand. The unwinding rate under these conditions is similar to the unwinding rate mediated by the E2 protein alone. when the E1 or E2 protein unwinds dsDNA independently, the unwinding rate of E1 is significantly faster than that of E2 (Figure 4F).

**Figure 4.**
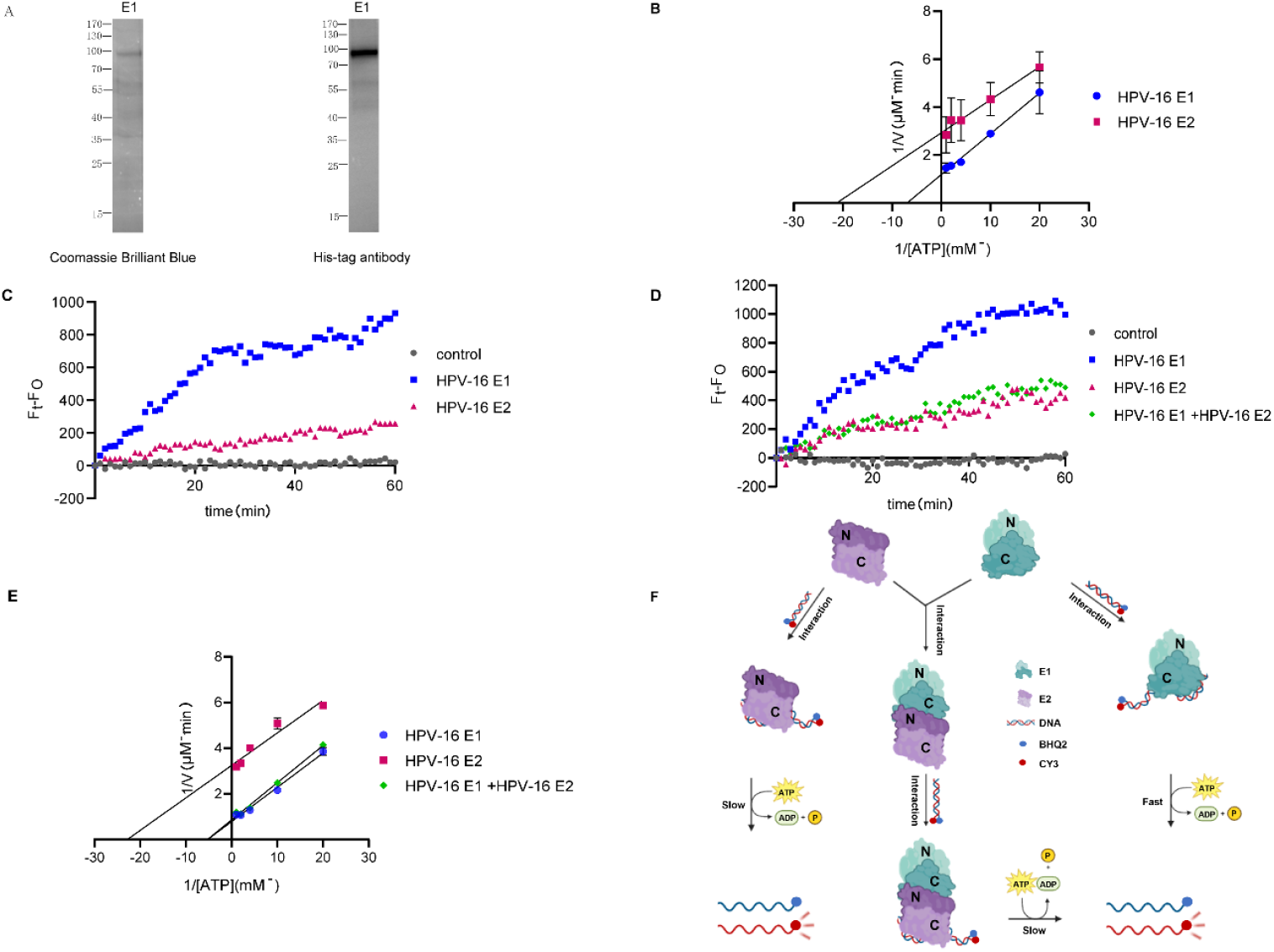
Comparison of enzymatic activities between E2 and E1 proteins and examination of E2’s effect on E1 activity. **(A)** SDS-PAGE analysis of purified E1 protein stained with Coomassie blue. Western blot analysis of purified E1 protein using an anti-His antibody. **(B)** Comparison of ATPase activity between 1 μM wild-type E2 and wild-type E1 proteins. **(C)** Real-time unwinding kinetics of 0.4 μM E2 and E1 proteins on fluorescently labeled 16-bp dsDNA. Buffer without protein served as a control. **(D)** Real-time unwinding kinetics of E1 protein alone or in the presence of E2 protein on fluorescently labeled 16-bp dsDNA. **(E)** ATPase activity of E2 protein alone and E1 protein with or without E2. ATP hydrolysis assays were performed at indicated ATP concentrations. Error bars represent the standard error of replicate measurements. **(F)** Schematic model of E2-mediated regulation of E1 enzymatic activity. The N-terminal domain of E1 (residues 1-143) is shown in light green, while its C-terminal domain is in dark green. The N-terminal domain of E2 (residues 1-245) is depicted in dark purple, and its C-terminal domain (residues 245-365) in light purple. “Slow” indicates slow unwinding of dsDNA, while “Fast” indicates rapid unwinding.

### E2 Inhibits E1 Helicase Activity Through Protein-Protein Interaction

To further validate whether E2 inhibits E1 helicase activity through direct interaction, we first confirmed the physical interaction between wild-type E2 and E1 proteins. Co-immunoprecipitation experiments demonstrated their interaction *in vitro* (Figure 5A). Molecular docking analysis revealed that the N-terminal domain of E2 (residues 1-245) binds to the C-terminal region of E1 (residues 143-645), with a specific hydrogen bond formed between E39 of E2 and N597 of E1 (Figure 5B), confirming our previous hypothesis (Figure 4F). To disrupt the E2-E1 protein interaction, the conserved residue E39 was mutated to alanine (Supplementary Figure S2), and an N-terminal truncation mutant of E2 (residues 1-245) was designed to further validate the E1-binding domain. Western blot analysis confirmed proper expression of both E39A and N-terminal truncation proteins (Supplementary Figure 5E). Subsequent co-immunoprecipitation experiments showed that while the N-terminal truncation maintained interaction with E1, the E39A mutation completely abolished this binding (Figure 5C), demonstrating that E2 interacts with E1 through its N-terminal domain (1-245) and that E39 is a critical residue for this interaction. Using FRET-based real-time helicase assays (Figure 5D), we observed that when the E39A mutation disrupted E2-E1 interaction, the unwinding activities of both proteins became additive. In contrast, the N-terminal truncation (1-245) and wild-type E2 maintained inhibition of E1 helicase activity, with unwinding kinetics similar to wild-type E2 (Figure 5D). These results conclusively demonstrate that E2 inhibits E1 helicase activity through direct protein-protein interaction.

**Figure 5.**
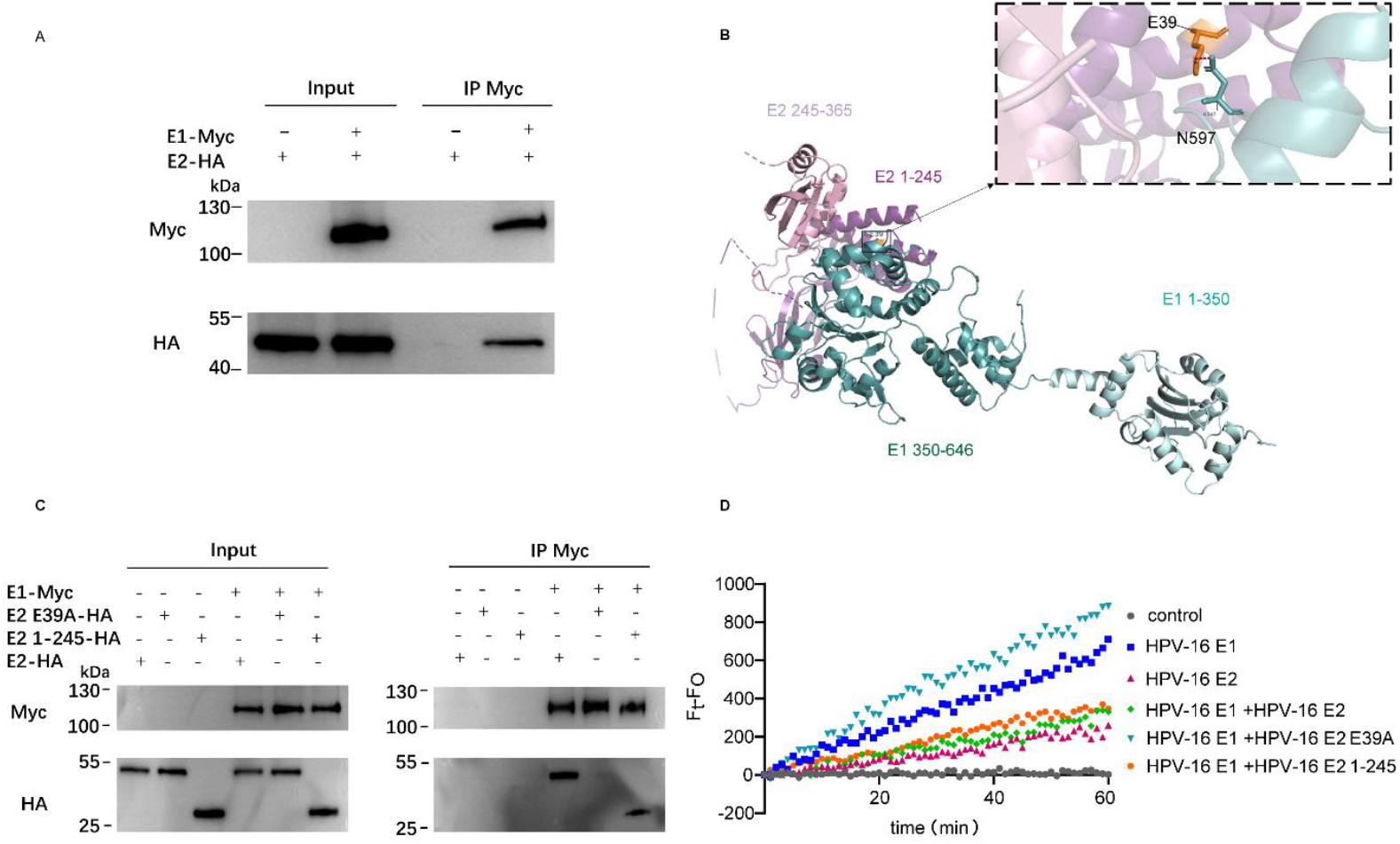
E2 inhibits E1 helicase activity through protein-protein interaction. **(A)** Interaction between E2 and E1 proteins *in vitro*. After purification of E2-HA and E1-Myc in *E. coli* BL21, co-immunoprecipitation (Co-IP) was performed, followed by Western blotting using anti-Myc or anti-HA antibodies. **(B)** HDOCK-based prediction of key interaction sites between E1 and E2 proteins. The N-terminal domain of E2 (residues 1-245) is shown in purple, while its C-terminal domain (245-365) is in pink. The N-terminal domain of E1 (1-350) is depicted in light green, and its C-terminal domain (350-646) in dark green. In the figure, E39 is colored in brown, N597 is shown in dark green, and hydrogen bonds are depicted in red. **(C)** Co-IP assay examining the interaction of E1 with wild-type (WT), mutant (E39A), and truncated E2 proteins (1-245), analyzed by Western blotting using anti-Myc or anti-HA antibodies. **(D)** Helicase assays measuring dsDNA unwinding by WT E2, E39A mutant, or truncated E2 (1-245) in the presence or absence of E1. Reactions contained 0.4 μM E2 (mutant or truncation) and E1, with 16-bp dsDNA as substrate.

## Discussion

This study provides compelling evidence through systematic biochemical and structural analyses that HPV 16 E2 protein possesses intrinsic ATPase and helicase activities. Mutagenesis studies identified critical residues (Y303 for ATPase activity; K299/Y303/K306 for helicase activity) that localize the enzymatic domain to the C-terminal region (245-365). This discovery challenges the conventional characterization of E2 as merely an auxiliary protein. While we have elucidated enzymatic features of E2 and its sophisticated regulatory interplay with E1 helicase, providing new perspectives on HPV DNA replication. Key unanswered questions include whether E2 maintains enzymatic activity in cellular and in vivo environments, and how mutations at critical residues affect HPV replication.

Particularly noteworthy is the finding that podophyllotoxin (PPT), an existing anti-HPV therapeutic, effectively inhibits E2 helicase activity through direct binding (IC50 = 0.1074 μM). This suggests its potential as an inhibitor targeting a critical viral protein. Compared to existing E6/E7-targeting inhibitors (e.g., Arenobufagin, MS-275, E7-PROTAC) (Schwartz et al,2022; Niu et al,2021; Jiang et al,2023), targeting E2 may block viral replication earlier and reduce the risk of genomic integration. However, the inhibitory effects of PPT on E2 and its anti-HPV activity in cells and in vivo require further investigation.

Research on the E1-E2 interaction has led to a groundbreaking discovery. Although enzymatic activity of E2 is weaker than E1, it exerts a dominant-negative effect through physical binding. HDOCK modeling and mutagenesis data reveal that N-terminal domain (1-245) of E2 binds helicase domain of E1, with E39 forming a crucial hydrogen bond (E39-N597). This interaction is characterized by two key features: 1) steric hindrance that prevents E1 from binding to the DNA substrate; and 2) reduction of the overall unwinding efficiency to the slower level characteristic of E2. This regulatory mechanism suggests a temporal control model for HPV replication: during initial stages, the potent helicase activity of E1 drives genomic amplification; as E2 expression increases, it modulates replication frequency by attenuating E1 activity while providing its own helicase function. Such fine-tuned regulation may prevent excessive replication that could trigger host defense mechanisms.

The discovery of E2 enzymatic activity and its functional interplay with E1 has opened new avenues for antiviral drug development. The inhibitory effect of podophyllotoxin particularly highlights the potential of targeting E2 helicase activity. Future studies should focus on: 1) the structural basis of E2 helicase function; 2) the physiological significance of this enzymatic activity in the viral life cycle; and 3) the development of specific E2 helicase inhibitors. These findings fundamentally expand our understanding of HPV replication mechanisms and provide a new framework for investigating auxiliary proteins in other viral systems.

## Methods

### Reagents

All reagents used in the experiments include: Anti-HA Tag Mouse Monoclonal Antibody (Selleck), Anti-MYC Tag Mouse Monoclonal Antibody (Selleck), Anti-His Tag Mouse Monoclonal Antibody (5C3) (Abbkine), ATP (CAS: 56-65-5, Solarbio), Podophyllotoxin (CAS: 518-28-5, Aladdin).

### Plasmid Construction and Mutagenesis

HPV 16 E1/E2 sequences were amplified using gene-specific primers and inserted into designated vectors via restriction enzyme digestion and ligation. Site-directed mutagenesis primers were designed based on the wild-type template sequence, targeting upstream and downstream regions of the desired mutation sites. Primer parameters were optimized using online tools (e.g., QuikChange® Site-Directed Mutagenesis Manual) to ensure matching Tm values and correct placement of mismatched bases. Mutants and truncations of E2 were generated through site-directed mutagenesis. All constructs were verified by sequencing.

### Recombinant Protein Expression and Purification

All recombinant proteins were expressed in *E. coli* BL21. Cells transformed with HPV 16-pET28a plasmids were cultured in Luria-Bertani medium containing kanamycin (100 μg/mL) at 37 °C until OD_600_ reached 0.8. Protein expression was induced with 0.4 mM isopropyl-β-D-thiogalactopyranoside (IPTG) at 16 °C for 24 hours. Cells were harvested, resuspended in lysis buffer (20 mM Tris-HCl, 500 mM NaCl, 0.5% Triton X-100, 20 mg/L RNase A, 20 mg/L DNase I, 10% glycerol, 1 mM DTT, protease inhibitors, pH 8.0), and lysed using a high-pressure homogenizer at 4 °C (1,300 Pa, 7 cycles). The lysate was centrifuged at 18,000 rpm for 45 minutes at 4 °C. His-tagged proteins were purified from the supernatant using Ni-NTA agarose beads with gentle inversion mixing for 4 hours at 4 °C. Discard the supernatant and transfer the beads to a binding column. Add wash buffer (20 mmol/L Tris-HCl, 500 mmol/L NaCl, 0.4 mmol/L DTT, 30 mmol/L imidazole, pH 8.0), mix well, and incubate on ice for 3 minutes. Then, drain the liquid from the column. Repeat the above operation. Continue washing the beads with a higher concentration of imidazole solution (20 mmol/L Tris-HCl, 500 mmol/L NaCl, 0.4 mmol/L DTT, 60 mmol/L imidazole, pH 8.0) until the effluent no longer turns blue upon addition of Coomassie Brilliant Blue G250 staining solution. After washing, completely drain the liquid from the beads. Add elution buffer (20 mmol/L Tris-HCl, 500 mmol/L NaCl, 500 mmol/L imidazole, 1 mmol/L DTT, 10% glycerol, pH 8.0), mix well, and incubate on ice for 5 minutes. Collect the liquid from the column. The eluate was dialyzed twice against buffer (100 mM NaCl, 20 mM Tris-HCl pH 8.0, 10% glycerol, 1 mM DTT) for 12 and 4 hours, respectively. Proteins were concentrated to approximately 0.8 mg/mL using 30/50 kDa cutoff concentrators (Solarbio), flash-frozen in liquid nitrogen, and stored at −80 °C.

### Protein Structure Prediction and Molecular Docking

HPV16 E1/E2 protein sequences were retrieved from http://www.ncbi.nlm.nih.gov/. Three-dimensional structures were predicted using https://alphafold3.org/. Protein interactions with small molecules, nucleic acids, or other proteins were predicted via http://hdock.phys.hust.edu.cn/ or AutoDock software. Resulting PDB files were visualized using PyMOL.

### ATPase Activity Assay

ATPase activity was measured using the QuantiChrom™ ATPase/GTPase Assay Kit (BioAssay Systems) to quantify inorganic phosphate released from ATP hydrolysis. Phosphate standards were prepared according to the manufacturer’s instructions. Reactions were performed in 96-well plates with 40 μL reaction volume per well. Wild-type or mutant HPV16 E proteins (0.4 μM) were incubated with varying ATP concentrations in buffer (20 mM Tris, 40 mM NaCl, 4 Mg(AcO)_2_, 0.5 mM EDTA, pH 7.5) at 37 °C for 1 hour. Dialysis buffer without protein served as the control. Reactions were terminated by adding 200 μL malachite green reagent, incubated for 30 minutes at room temperature, and absorbance was measured at 620 nm using a microplate reader. Reaction velocities and ATP concentrations fit the Michaelis-Menten equation: 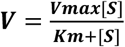

### Helicase Activity Assay

Helicase activity was assessed using a FRET-based dsDNA unwinding assay. Complementary DNA oligonucleotides labeled with Cy3 or BHQ2 were synthesized and dissolved in annealing buffer (20 mM Tris-NaCl, 2 mM MgCl_2_, 0.2 mM DTT, 100 mM KCl, pH 7.5) to 10 μM. dsDNA was prepared by mixing equimolar complementary strands to a final concentration of 500 nM, heating to 95 °C for 5 minutes, and slowly cooling to room temperature. Unwinding reactions contained 0.4 μM purified HPV16 E2 protein, 50 nM dsDNA substrate, 5 mM ATP, and 500 nM unlabeled competitor DNA (5’-GCGTCTTTACGGTGCT-3’)in reaction buffer (20 mM Tris-HCl pH 7.0, 10 mM NaCl, 0.1 mg/mL BSA, 5 mM MgCl_2_, 2 mM DTT) in a total volume of 100 μL. Reactions were transferred to white 96-well plates, covered with foil, and fluorescence was measured every minute for 1 hour using a microplate reader (excitation: 550 nm, emission: 620 nm, cutoff: 610 nm).

### Co-Immunoprecipitation (Co-IP)

E1 and E2 Proteins were mixed at a 1:1 molar ratio in binding buffer (0.5% Triton X-100 in PBS) and incubated with rotation at 4 °C for 2 hours. MYC antibody was added and incubated for another 2 hours, followed by addition of 20 μL Protein A/G beads and further rotation for 2 hours. Beads were pelleted by centrifugation at 12,000 rpm for 2 minutes at 4 °C, washed three times with wash buffer (20 mM Tris–HCl, 300 mM NaCl, 0.1 mM EDTA, pH 8.0), resuspended in 1× protein loading buffer, boiled for 10 minutes, and centrifuged. The supernatant was collected for Western blot analysis.

### Bio-Layer Interferometry (BLI)

Bio-Layer Interferometry (BLI) was employed to quantitatively characterize the binding interaction between HPV16 E2 protein and PPT. This technique utilizes fiber optic biosensors to monitor changes in the optical thickness of the sensor layer resulting from biomolecular binding events (Murali et al, 2022; Miczi et al,2021; Durous et al, 2024; Boclinville et al, 2024). In this system, analytes interacting with ligands immobilized on the sensor surface form a monolayer, leading to proportional shifts in the interference spectrum of reflected light. Analyses were performed on a ForteBio Octet RED96 system equipped with dedicated data acquisition and analysis software (Pall ForteBio Corp., CA, USA).All steps were carried out at 25°C with a total working volume of 200 μL for samples and buffers, loaded in the instrument-specific 96-well black plates. Prior to measurement, Ni-NTA sensors were pre-wet in protein storage buffer (100 mM NaCl, 20 mM Tris-HCl pH 8.0, 10% glycerol, and 1 mM DTT) for 10 minutes. The loading procedure consisted of three steps: (1) baseline (60 s), (2) loading (60 s), and (3) baseline (60 s). After loading, the Ni-NTA biosensors bound with E2 protein and control sensors were equilibrated in assay buffer before being transferred into wells containing different concentrations of podophyllotoxin for binding measurement.TBS-T (137 mM NaCl, 2.7 mM KCl, 25 mM Tris, 0.05% Tween-20) served as the assay buffer, with podophyllotoxin concentrations tested at 50, 100, 200, and 400 μM. The binding assay protocol comprised five steps: (1) baseline (300 s), (2) protein association (120 s), (3) baseline (60 s), (4) small molecule association (60 s), and (5) dissociation (60 s). Finally, kinetic parameters (kon and koff) and the affinity constant (KD) were obtained using a double-referencing subtraction method (sample and sensor reference) based on a 1:1 binding model, processed with the data analysis software (Octet BLI Analysis 12.2).

### IC_50_ Determination of PPT as an E2 Helicase Inhibitor

PPT was serially diluted in DMSO to concentrations ranging from 0.1 mM to 0.0001 mM. E2 protein (0.4μM) was pre-incubated with DMSO or PPT for 20 minutes, followed by addition of reaction buffer containing 50 nM dsDNA substrate, 4 mM ATP, and 500 nM competitor DNA. Reactions were carried out in white 96-well plates (100 μL total volume) covered with foil. After 1 hour, fluorescence was measured (excitation: 550 nm, emission: 620 nm, cutoff: 610 nm). Experiments were performed in triplicate. Inhibition rate was calculated as:

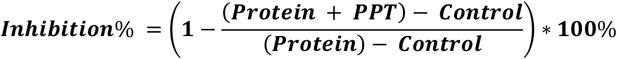

Data were fitted using GraphPad Prism 8 with the “log(inhibitor) vs. response – Variable slope (four parameters)” model.

## Data availability

This study includes no data deposited in external repositories.

## Author contributions

**Ping Xu**: Conceptualization, Methodology, Formal analysis, Investigation, Validation, Writing - original draft, Writing - review & editing. **Shuning Cai**: Methodology, Investigation, Writing - review & editing. **Lihong Zhang:** Writing – Review & Editing. **Yueqi Wu:**Writing – Review & Editing. **Shan Xu**: Conceptualization, Resources, Supervision, Funding acquisition, Writing - review & editing. **Yigang Tong**: Conceptualization, Resources, Supervision, Funding acquisition, Writing - review & editing. **Ke Xu**: Funding acquisition, Writing - review & editing.

## Acknowledgements

This work was supported by Fundamental Research Funds for the Central Universities (buctrc202230), Capital Health Research and Development of Special Fund (2024-1G-4421), National Natural Science Foundation of China (82204230) and the Shandong Provincial Natural Science Foundation (ZR2022QH054).

## Supplementary Data

Supplementary data is available in the “Supplementary Data.docx” document.

## References

1. Bruni L, Albero G, Rowley J, Alemany L, Arbyn M, Giuliano AR, Markowitz LE, Broutet N, Taylor M. (2023). Global and regional estimates of genital human papillomavirus prevalence among men: a systematic review and meta-analysis. Lancet Glob Health, 11(9): e1345–e1362.

2. Deng SJ, Pearce KH, Dixon EP, Hartley KA, Stanley TB, Lobe DC, Garvey EP, Kost TA, Petty RL, Rocque WJ et al. (2004). Identification of peptides that inhibit the DNA binding, trans-activator, and DNA replication functions of the human papillomavirus type 11 E2 protein. J Virol, 78(5): 2637–2641.

3. Sung H, Ferlay J, Siegel RL, Laversanne M, Soerjomataram I, Jemal A, Bray F. (2021). Global Cancer Statistics 2020: GLOBOCAN Estimates of Incidence and Mortality Worldwide for 36 Cancers in 185 Countries. CA Cancer J Clin, 71(3): 209–249.

4. McBride AA. (2022). Human papillomaviruses: diversity, infection and host interactions. Nat Rev Microbiol, 20(2): 95–108.

5. Arbyn M, Weiderpass E, Bruni L, de Sanjosé S, Saraiya M, Ferlay J, Bray F. (2020). Estimates of incidence and mortality of cervical cancer in 2018: a worldwide analysis. Lancet Glob Health, 8(2): e191–e203.

6. Pešut E, Ðukić A, Lulić L, Skelin J, Šimić I, Milutin Gašperov N, Tomaić V, Sabol I, Grce M. (2021). Human Papillomaviruses-Associated Cancers: An Update of Current Knowledge. Viruses, 13(11): 2234.

7. Serrano B, Alemany L, Tous S, Bruni L, Clifford GM, Weiss T, Bosch FX, de Sanjosé S. (2012). Potential impact of a nine-valent vaccine in human papillomavirus related cervical disease. Infect Agent Cancer, 7(1): 38.

8. D’Souza MJ, Hundi GK, Dandekeri S, Jayaraman J. (2023). Intralesional Measles, Mumps and Rubella Vaccine versus Formic Acid Puncture in the Treatment of Common Warts: A Prospective Randomised Study. Indian J Dermatol, 68(4): 486.

9. Haley CT, Mui UN, Vangipuram R, Rady PL, Tyring SK. (2019). Human oncoviruses: Mucocutaneous manifestations, pathogenesis, therapeutics, and prevention: Papillomaviruses and Merkel cell polyomavirus. J Am Acad Dermatol, 81(1): 1–21.

10. Muñoz N, Bosch FX, de Sanjosé S, Herrero R, Castellsagué X, Shah KV, Snijders PJ, Meijer CJ; International Agency for Research on Cancer Multicenter Cervical Cancer Study Group. (2003). Epidemiologic classification of human papillomavirus types associated with cervical cancer. N Engl J Med, 348(6): 518–527.

11. Al-Awadhi R, Al-Mutairi N, Albatineh AN, Chehadeh W. (2019). Association of HPV genotypes with external anogenital warts: a cross sectional study. BMC Infect Dis, 19(1): 375.

12. Koster S, Gurumurthy RK, Kumar N, Prakash PG, Dhanraj J, Bayer S, Berger H, Kurian SM, Drabkina M, Mollenkopf HJ et al. (2022). Modelling Chlamydia and HPV co-infection in patient-derived ectocervix organoids reveals distinct cellular reprogramming. Nat Commun, 13(1): 1030.

13. Arroyo Mühr LS, Gini A, Yilmaz E, et al. (2024) Concomitant human papillomavirus (HPV) vaccination and screening for elimination of HPV and cervical cancer. Nat Commun 15: 3679.

14. Whop LJ, Cunningham J, Garvey G, Condon JR. (2019). Towards global elimination of cervical cancer in all groups of women. Lancet Oncol, 20(5): e238.

15. Billingsley CL, Chintala S, Katzenellenbogen RA. (2022). Post-Transcriptional Gene Regulation by HPV 16E6 and Its Host Protein Partners. Viruses, 14(7): 148.

16. Schiffman M, Castle PE, Jeronimo J, Rodriguez AC, Wacholder S. (2007). Human papillomavirus and cervical cancer. Lancet, 370(9590): 890–907.

17. Meng Q, Zhang Y, Sun H, Yang X, Hao S, Liu B, Zhou H, Wang Y, Xu ZX. (2024). Human papillomavirus-16 E6 activates the pentose phosphate pathway to promote cervical cancer cell proliferation by inhibiting G6PD lactylation. Redox Biol, 71: 103108.

18. Nelson CW, Mirabello L. (2023). Human papillomavirus genomics: Understanding carcinogenicity. Tumour Virus Res, 15: 200258.

19. Hebner CM, Laimins LA. (2006). Human papillomaviruses: basic mechanisms of pathogenesis and oncogenicity. Rev Med Virol, 16(2): 83–97.

20. Sakakibara N, Mitra R, McBride AA. (2011). The papillomavirus E1 helicase activates a cellular DNA damage response in viral replication foci. J Virol, 85(17): 8981–8995.

21. Nakahara T, Tanaka K, Ohno S, Egawa N, Yugawa T, Kiyono T. (2015). Activation of NF-κB by human papillomavirus 16 E1 limits E1-dependent viral replication through degradation of E1. J Virol, 89(9): 5040–5059.

22. Schuck S, Stenlund A. (2011). Mechanistic analysis of local ori melting and helicase assembly by the papillomavirus E1 protein. Mol Cell, 43(5): 776–787.

23. Arbyn M, Weiderpass E, Bruni L, de Sanjosé S, Saraiya M, Ferlay J, Bray F. (2020). Estimates of incidence and mortality of cervical cancer in 2018: a worldwide analysis. Lancet Glob Health, 8(2): e191–e203.

24. Yilmaz G, Biswas-Fiss EE, Biswas SB (2023) Sequence-Dependent Interaction of the Human Papillomavirus E2 Protein with the DNA Elements on Its DNA Replication Origin. Int J Mol Sci 24: 6555.

25. Sedman J, Stenlund A. (1998). The papillomavirus E1 protein forms a DNA-dependent hexameric complex with ATPase and DNA helicase activities. J Virol, 72(8): 6893–6897.

26. Longworth MS, Laimins LA. (2004). Pathogenesis of human papillomaviruses in differentiating epithelia. Microbiol Mol Biol Rev, 68(2): 362–372.

27. Chen G, Stenlund A. (2002). Sequential and ordered assembly of E1 initiator complexes on the papillomavirus origin of DNA replication generates progressive structural changes related to melting. Mol Cell Biol, 22(21): 7712–7720.

28. Baedyananda F, Sasivimolrattana T, Chaiwongkot A, Varadarajan S, Bhattarakosol P. (2022). Role of HPV16 E1 in cervical carcinogenesis. Front Cell Infect Microbiol, 12: 955847.

29. Orav M, Gagnon D, Archambault J. (2019). Interaction of the Human Papillomavirus E1 Helicase with UAF1-USP1 Promotes Unidirectional Theta Replication of Viral Genomes. mBio, 10(2): e00152–19.

30. Bandyopadhyay S, Raney KD, Liu Y, Hermonat PL. (2008). AAV-2 Rep78 and HPV-16 E1 interact in vitro, modulating their ATPase activity. Biochemistry, 47(2): 845–856.

31. Hughes FJ, Romanos MA. (1993). E1 protein of human papillomavirus is a DNA helicase/ATPase. Nucleic Acids Res, 21(25): 5817–5823.

32. Rana A, Yilmaz G, Biswas-Fiss EE, Biswas S. (2025). Mechanisms of Viral DNA Replication of Human Papillomavirus: E2 Protein-Dependent Recruitment of E1 DNA Helicase to the Origin of DNA Replication. Int J Mol Sci, 26(9): 4333.

33. Deng W, Lin BY, Jin G, Wheeler CG, Ma T, Harper JW, Broker TR, Chow LT. (2004). Cyclin/CDK regulates the nucleocytoplasmic localization of the human papillomavirus E1 DNA helicase. J Virol, 78(24): 13954–13965.

34. Schwartz J, Harris C, Pietryga J, Zheng H, Kumar P, Visheratina A, Kotov NA, Major B, Avery P, Ercius P, et al.(2022). Real-time 3D analysis during electron tomography using tomviz. Nat Commun, 13(1): 4458.

35. Niu C, Leavitt LS, Lin Z, Paguigan ND, Sun L, Zhang J, Torres JP, Raghuraman S, Chase K, Cadeddu R et al. (2021). Neuroactive Type-A γ-Aminobutyric Acid Receptor Allosteric Modulator Steroids from the Hypobranchial Gland of Marine Mollusk, Conus geographus. J Med Chem, 64(10): 7033–7043.

36. Jiang B, Weinstock DM, Donovan KA, Sun HW, Wolfe A, Amaka S, Donaldson NL, Wu G, Jiang Y, Wilcox RA et al. (2023). ITK degradation to block T cell receptor signaling and overcome therapeutic resistance in T cell lymphomas. Cell Chem Biol, 30(4): 383–393.e6.

37. Hou SY, Wu SY, Chiang CM. (2002). Transcriptional activity among high and low risk human papillomavirus E2 proteins correlates with E2 DNA binding. J Biol Chem, 277(47): 45619–45629.

38. McBride AA. (2013). The papillomavirus E2 proteins. Virology, 445(1-2): 57–79.

39. Doorbar J, Egawa N, Griffin H, Kranjec C, Murakami I. (2015). Human papillomavirus molecular biology and disease association. Rev Med Virol, 25(Suppl 1): 2–23.

40. Doorbar J, Quint W, Banks L, Bravo IG, Stoler M, Broker TR, Stanley MA. (2012). The biology and life-cycle of human papillomaviruses. Vaccine, 30(Suppl 5): F55–F70

41. Bray F, Laversanne M, Sung H, Ferlay J, Siegel RL, Soerjomataram I, Jemal A. (2024). Global cancer statistics 2022: GLOBOCAN estimates of incidence and mortality worldwide for 36 cancers in 185 countries. CA Cancer J Clin, 74(3): 229–263.

42. Brisson M, Kim JJ, Canfell K, Drolet M, Gingras G, Burger EA, Martin D, Simms KT, Bénard É, Boily MC et al, (2020). Impact of HPV vaccination and cervical screening on cervical cancer elimination: a comparative modelling analysis in 78 low-income and lower-middle-income countries. Lancet, 395(10224): 575–590.

43. Drolet M, Laprise JF, Martin D, Jit M, Bénard É, Gingras G, Boily MC, Alary M, Baussano I, Hutubessy R, Brisson M. (2021). Optimal human papillomavirus vaccination strategies to prevent cervical cancer in low-income and middle-income countries in the context of limited resources: a mathematical modelling analysis. Lancet Infect Dis, 21(11): 1598–1610.

44. Kollipara R, Ekhlassi E, Downing C, Guidry J, Lee M, Tyring SK. (2015). Advancements in Pharmacotherapy for Noncancerous Manifestations of HPV. J Clin Med, 4(5): 832–846.

45. Komericki P, Akkilic-Materna M, Strimitzer T, Aberer W. (2011). Efficacy and safety of imiquimod versus podophyllotoxin in the treatment of anogenital warts. Sex Transm Dis, 38(3): 216–218.

46. Gilson R, Nugent D, Bennett K, Doré CJ, Murray ML, Meadows J, Haddow LJ, Lacey C, Sandmann F, Jit M, Soldan K et al. (2020). Imiquimod versus podophyllotoxin, with and without human papillomavirus vaccine, for anogenital warts: the HIPvac factorial RCT. Health Technol Assess, 24(47): 1–86.

47. Murray ML, Meadows J, Doré CJ, Copas AJ, Haddow LJ, Lacey C, Jit M, Soldan K, Bennett K, Tetlow M et al. (2018). Human papillomavirus infection: protocol for a randomised controlled trial of imiquimod cream (5%) versus podophyllotoxin cream (0.15%), in combination with quadrivalent human papillomavirus or control vaccination in the treatment and prevention of recurrence of anogenital warts (HIPvac trial). BMC Med Res Methodol, 18(1): 125.

48. Nicolaidou E, Kanelleas A, Nikolakopoulos S, Bezrodnii G, Nearchou E, Gerodimou M, Papadopoulou-Skordou E, Paparizos V, Rigopoulos D. (2021). A short, 8-week course of imiquimod 5% cream versus podophyllotoxin in the treatment of anogenital warts: A retrospective comparative cohort study. Indian J Dermatol Venereol Leprol, 87(5): 666–670.

49. Murali S, Rustandi RR, Zheng X, Payne A, Shang L (2022) Applications of Surface Plasmon Resonance and Biolayer Interferometry for Virus-Ligand Binding. Viruses 14: 717.

50. Miczi M, Diós Á, Bozóki B, Tőzsér J, Mótyán JA. (2021). Development of a Bio-Layer Interferometry-Based Protease Assay Using HIV-1 Protease as a Model. Viruses, 13(6): 1183.

51. Durous L, Géraudie S, Andersen L, et al. (2024) Rapid identity testing of antibody-based hot targeted radionuclide therapies by bio-layer interferometry. J Pharm Biomed Anal 246: 116227.

52. Boclinville A, Vandevenne M, Ambroggio E, et al. (2024) Interaction studies between human papillomavirus virus-like particles and laminin 332 by affinity capillary electrophoresis assisted by bio-layer interferometry. Talanta 270: 125602.

